# Dataset Decay: the problem of sequential analyses on open datasets

**DOI:** 10.1101/801696

**Authors:** William Hedley Thompson, Jessey Wright, Patrick G Bissett, Russell A Poldrack

## Abstract

Open data has two principal uses: (i) to reproduce original findings and (ii) to allow researchers to ask new questions with existing data. The latter enables discoveries by allowing a more diverse set of viewpoints and hypotheses to approach the data, which is self-evidently advantageous for the progress of science. However, if many researchers reuse the same dataset, multiple statistical testing may increase false positives in the literature. Current practice suggests that the number of tests to be corrected is the number of simultaneous tests performed by a researcher. Here we demonstrate that sequential hypothesis testing on the same dataset by multiple researchers can inflate error rates. This finding is troubling because, as more researchers embrace an open dataset, the likelihood of false positives (i.e. type I errors) will increase. Thus, we should expect a dataset’s utility for discovering new true relations between variables to decay. We consider several sequential correction procedures. These solutions can reduce the number of false positives but, at the same time, can prompt undesired challenges to open data (e.g. incentivising restricted access).

## Introduction

In recent years, there has been a push to increase the adoption of open research practices, which includes making scientific datasets accessible (Nosek et al. 2015). Open data allow researchers to both reproduce published analyses and ask new questions of the existing data (Molloy 2011; Pisani et al. 2016). The value attributed to the latter is that it makes discoveries and the advancement of knowledge more efficient. After all, data often can be useful for investigating and discovering phenomena beyond its initial purpose. The proliferation and use of open data will increase over time as funders mandate and reward data sharing and open research practices (McKiernan et al. 2016).

While open data undoubtedly provides these benefits, a problem emerges regarding multiple hypothesis testing on a single dataset. At present, researchers reusing data generally do not take into account the previous studies that have performed tests on the dataset; instead, they only correct for the number of statistical tests that they perform. We will show that multiple reuses of the same dataset will compound type I error rates just as if the multiple tests were performed as part of a single analysis.

In statistics, a distinction is made between *simultaneous* and *sequential* correction procedures when correcting for multiple tests. Simultaneous procedures correct for all tests at once, while sequential procedures correct for the latest in a non-simultaneous series of tests. Several solutions have been proposed to address multiple sequential analyses, namely *α-spending* and *α-investing* procedures (Foster and Stine 2008; Aharoni and Rosset 2014). Here we will also propose a third, *α-debt*, which does not maintain a constant false positive rate but allows it to grow controllably.

Sequential correction procedures are harder to implement than simultaneous procedures as they require keeping track of the total number of tests that have been performed by others. Further, in order to ensure data is still shared, the sequential correction procedures should not be antagonistic with current data sharing incentives and infrastructure. Thus, we have identified several desiderata regarding open data and multiple hypothesis testing:

### Sharing incentive

Data producers should be able to share their data without negatively impacting their initial statistical tests. Otherwise, this reduces the incentive to share data.

### Open access

Minimal to no restrictions should be placed on accessing open data, other than those necessary to protect the confidentiality of human subjects. Otherwise, the data are no longer open.

### Stable false positive rate

The false positive rate (i.e. type I error) should not increase due to reusing open data. Otherwise, scientific results become less reliable with each reuse.

We will show that obtaining all three of these desiderata is not possible. We will demonstrate below that the current practice of ignoring sequential tests leads to an increased false positive rate in the scientific literature. Further, we show that sequentially correcting for data reuse can reduce the number of false positives compared to current practice. However, all the proposals considered here must still compromise (to some degree) on one of the above desiderata.

## Results

### Families of tests through time

Procedures to correct for multiple statistical tests predated open data as promoted today. These procedures were designed for situations in which a researcher performs multiple statistical tests within the same experiment. In general, statistical decisions involve a trade-off between the rate of false positives (type I errors) and the rate of false negatives (type II errors) (Ryan 1962; Hochberg and Tamhane 1987). These error rates can relate to an individual statistical test (Wilson 1962) or an entire experiment (Ryan 1959, 1962). Typically, error rates are considered for neither of these two extremes but rather for a *family* of tests, a set which includes some related statistical tests. Unfortunately, the term family has been challenging to precisely define, and only guidelines – often containing additional imprecise terminology – exist (e.g. Cox 1965; Miller 1981; Hochberg and Tamhane 1987; Hancock and Klockars 1996). Generally, tests are considered part of a family when: (i) multiple variables are being tested with no definitive hypothesis, or (ii) multiple prespecified tests together help support the same or associated research questions (Hochberg and Tamhane 1987; Hancock and Klockars 1996).

The crucial question for the present purpose is whether the reuse of data constitutes a new family of tests. If sequential analyses create a new family of tests, then there is no need to perform a sequential correction procedure in order to maintain control over familywise error. Alternatively, if a new family has not been created simply by reusing data, then we need to consider sequential correction procedures.

There are two ways in which sequential tests with open data differ from simultaneous tests (where correction is needed): a time lag between tests and different individuals performing the tests. Neither of these two properties is sufficient to justify the emergence of a new family of tests. First, the temporal displacement of statistical tests can not be considered sufficient reason for creating a new family of statistical tests, as the speed with which a researcher analyzes a dataset is not relevant to the need to control for multiple statistical tests. If it were, then a simple correction procedure would be to wait a specified length of time before performing the next statistical test. Second, it should not matter who performs the tests; otherwise, one could correct for multiple tests by crowd-sourcing the analysis. Thus if we were to decide that either of the two differentiating properties of sequential tests on open data creates a new family, undesirable procedures would be allowable. To prevent this, statistical tests on open data, which can be run by different people, and at different times, can be part of the same family of tests. Since they can be in the same family, sequential tests on open data need to consider correction procedures to control the rate of false positives across the family.

We have demonstrated the possibility that families of tests can belong to sequential analyses. However, in practice, when does this occur? Due to the fuzzy nature of “family”, we propose a simple rule-of-thumb: if the sequential tests would be considered within the same family if performed simultaneously, then they are part of the same family in sequential tests. Applying this rule indicates that many sequential tests should be considered part of the same family when reusing open data (see Supplementary Material for examples of sequential families). We therefore suggest that researchers should apply corrections for multiple tests when reusing data or provide a justification for the lack of such corrections (as they would need to in the case of simultaneous tests belonging to different families).

### The consequence of not taking multiple sequential testing seriously

In this section, we consider the consequences of uncorrected sequential testing and several procedures to correct for them. We start with a simulation to test the false positive rate of the different sequential correction procedures by performing 100 sequential univariate tests where the simulated covariance between all variables was 0 (see Methods for additional details). The simulations ran for 1,000 iterations, and the familywise error was calculated using a two-tailed statistical significance threshold of p < 0.05.

We first consider what happens when the sequential tests are uncorrected. Unsurprisingly, the results are identical to not correcting for simultaneous tests (Figure 1A). There will almost always be at least one false positive any time one performs 100 sequential analyses with this simulation. This rate of false positives is dramatically above the desired familywise error rate of at least one false positive in 5% of the simulation’s iterations. Uncorrected sequential tests will lead to more false positives.

**Figure 1:**
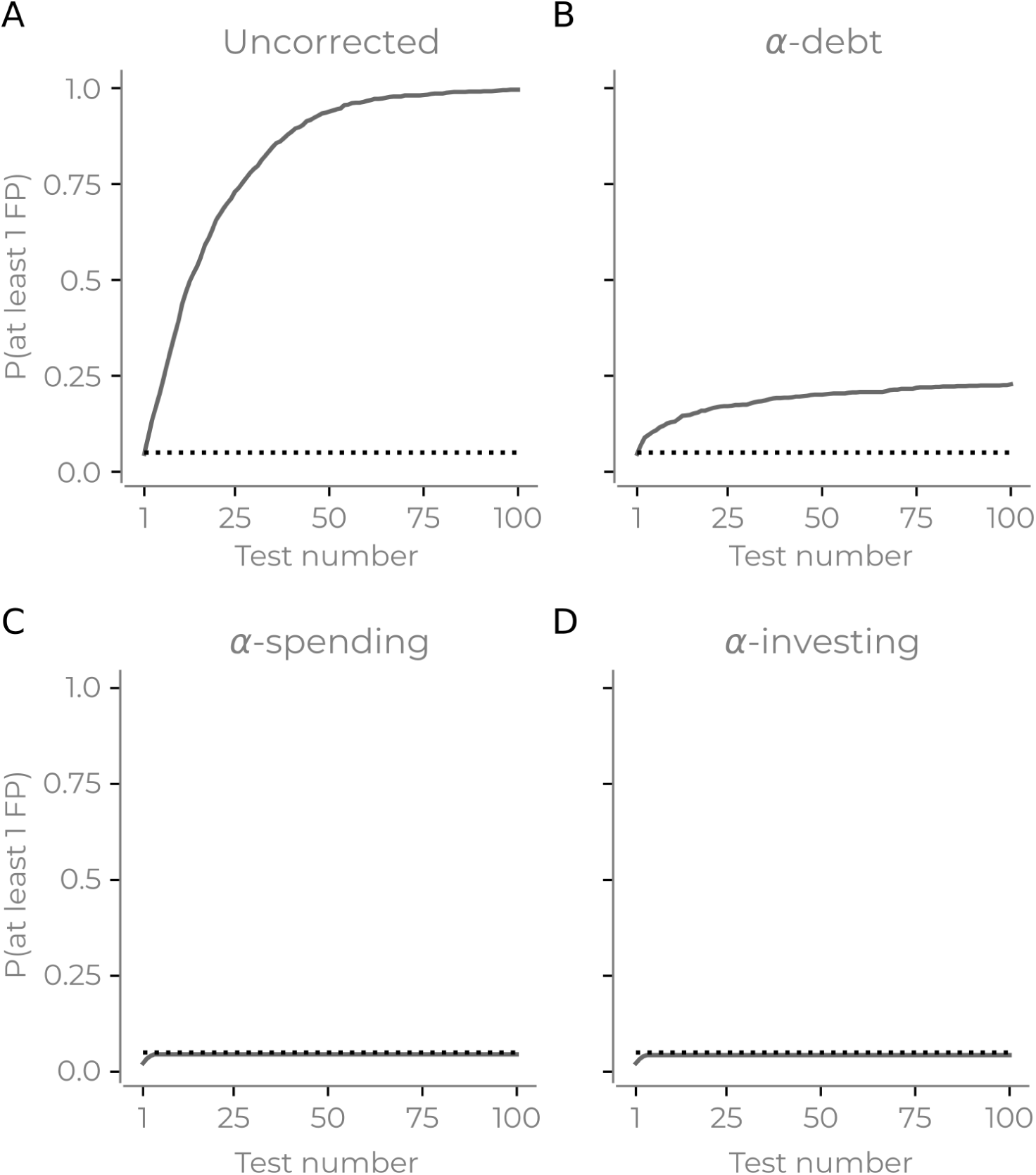
Simulation results showing the probability of there being at least one false positive as the number of statistical tests increases. Each panel shows different correction procedures: (A) uncorrected; (B) *α*-debt; (C) *α*-spending; (D) *α*-investing. Dotted line indicates 0.05.

The first sequential procedure we consider is *α-debt*. For the *i*th sequential test, this procedure considers there to be *i* simultaneous tests that should be corrected. This procedure effectively performs a Bonferroni correction – i.e. the threshold of statistical significance becomes 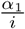 where *α*_1_ is the first statistical threshold (here 0.05). Thus, on the first test *α*_1_ = 0.05, then on the second sequential test *α*_2_ = 0.025, *α*_3_ = 0.0167, and so on. While each sequential test is effectively a Bonferroni correction considering all previous tests, this does not retroactively change the inference of any previous statistical tests. When a new test is performed, the previous test’s *α* is now too lenient considering all the tests that have been performed. Thus, when considering all tests together, the false positive rate will increase, accumulating a false positive “debt”. This debt entails that method does not ensure the type I error rate remains under a specific value, instead allows it to controllably increase under a “debt ceiling” with each sequential test (the debt ceiling is the sum of all *α*_1_ to *α*_*t*_ at *t*). The rate of the increase in debt always decreases. Both the increase in false positives and the decrease in debt increase were confirmed in the simulations (Figure 1B). Finally, the method can mathematically ensure that the type II error (i.e. the false negative rate) is equal to or better than simultaneous correction with Bonferroni (See Methods).

The next two procedures we consider have previously been suggested in the literature *α*-spending and *α*-investing (Foster and Stine 2008; Aharoni and Rosset 2014). The first has a total amount of “*α* wealth”, and the sum of all the statistical thresholds for all sequential tests can never exceed this amount (i.e. if the alpha wealth is 0.05 then the sum of all thresholds on sequential tests must be less than 0.05). Here, for each sequential test, we spend half the remaining wealth (i.e. *α*_1_ is 0.025, *α*_2_ is 0.0125 and so on). In the simulations, the sequential tests limit the probability of there being at least one false positive to less than 0.05 (Figure 1C). Finally, *α*-investing allows for the significance threshold to increase or decrease as researchers perform additional tests. Again there is a concept of *α*-wealth. If a test rejects the null hypothesis, there is an increase in the remaining *α*-wealth that future tests can use and, if the reverse occurs, the remaining *α*-wealth decreases (see methods). *α*-investing ensures control of the false discovery rate at an assigned level. Here we invest 50% of the remaining wealth for each statistical test. In the simulations, this method also remains under 0.05 familywise error rate in the simulations as the sequential tests increase. (Figure 1D).

The main conclusion from this set of simulations is that the current practice of not correcting for open data reuse results in a substantial increase in the number of false positives presented in the literature.

### Sensitivity to the order of sequential tests

The previous simulation did not consider any true positives in the data (i.e. cases where we should reject the null hypothesis). Since the statistical threshold for significance changes as the number of sequential tests increases, it becomes crucial to evaluate the sensitivity of each method to both type I and type II errors in regards to the order of sequential tests. Thus, we simulated true positives (between 1-10) where the covariance of these variables and the dependent variable were set to *p* (*p* ranged between 0 and 1). Further, *λ* controls the sequential test order determining the probability that a test was a true positive. When *λ* is positive, it entails a higher likelihood that earlier tests will be one of the true positives (and vice versa when *λ* was negative; see methods). All other parameters are the same as the previous simulation. Simultaneous correction procedures (Bonferroni and FDR) of all 100 tests were also included to contrast the different sequential procedures to these methods.

The results reveal that the order of the tests is pivotal for sequential correction procedures. Unsurprisingly, the uncorrected and simultaneous correction procedures do not depend on the sequential order of tests (Figure 2ABC). The sequential correction procedures all increased their true positive rate (i.e. less type II errors) when the true positives were earlier in the analysis order (Figure 2A). We also observe that *α*-debt had the highest true positive rate of the sequential procedures and, when the true positives were later in the test sequence, performed on par with Bonferroni but when the true positives were earlier, outperformed Bonferroni. *α*-investing and *α*-spending cannot give such assurances when the true positives are later in the analysis sequence (i.e. *λ* is negative) there is less sensitivity to true positives (i.e. type II errors). *α*-debt is more sensitive to true positives compared to *α*-spending because the threshold for the *m*h sequential test decreases linearly in *α*-debt and exponentially in *α*-spending. This results in a more lenient statistical threshold for *α*-debt in later sequential tests.

**Figure 2:**
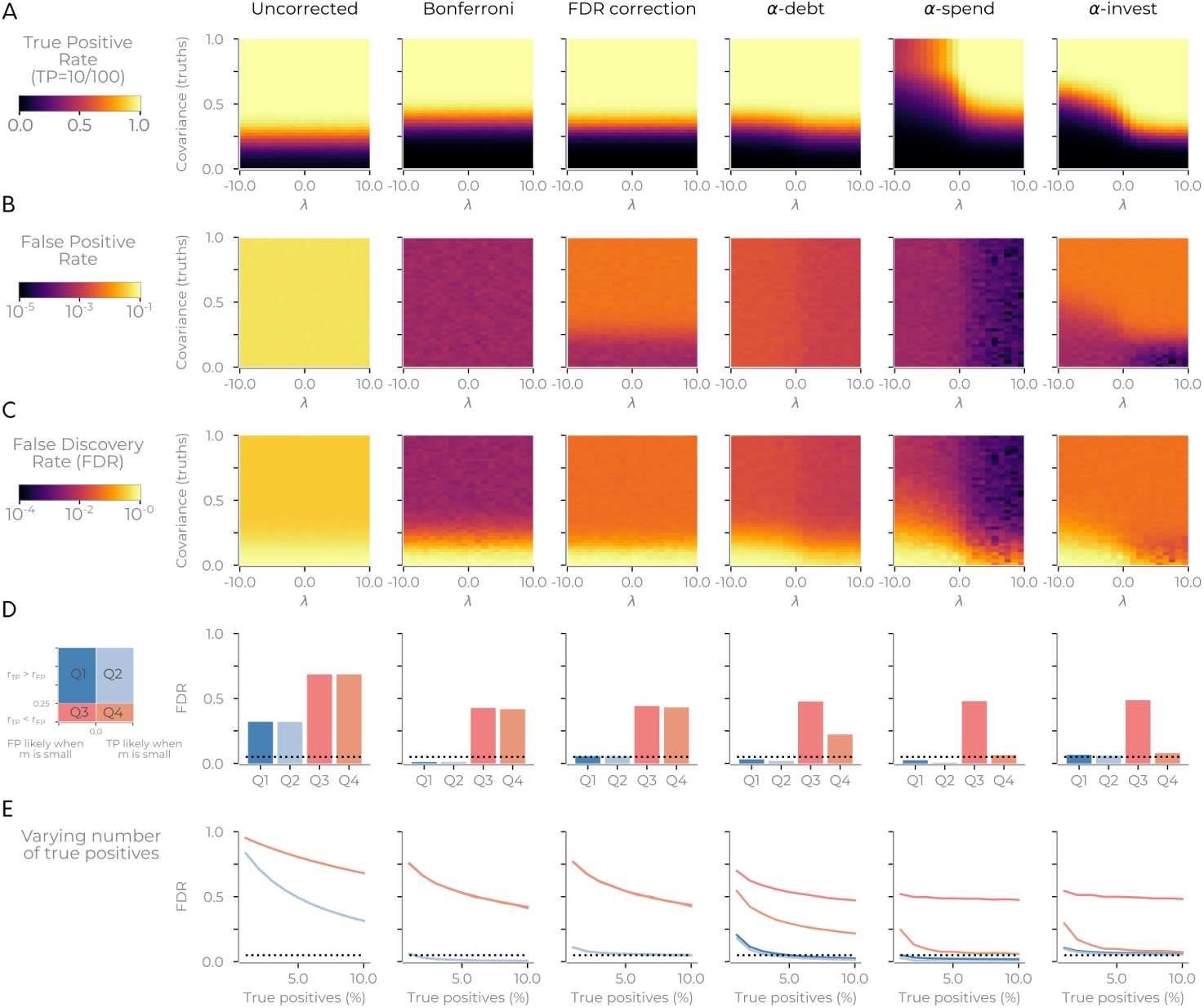
Results of simulations showing that the order of sequential tests can impact true positive sensitivity. (A) The true positive rate after 100 tests for different sequential correction procedures (for 1,000 iterations). Each procedure shows the effect of simulation parameters *λ* (when positive, it increases the probability of the true positives being an earlier test) and the simulated covariance of the true positives. The results showed simulations when there were ten true positives in the data. (B) Same as A, but shows the false positive rate. (C) Same as A, but shows the false discovery rate. (D) The panels in C split into four quadrants. The first split was the probability of true positives being an earlier test (Q2, Q4, *λ* > 0) and later tests (Q1, Q3, *λ* < 0). The second split was the covariance of the correlated variables (Q1, Q2, > 0.25; Q3, Q4 < 0.25). Panels show the average FDR for various quadrants from D. (E) Same as D but showing the varying number of true positives that existed in the simulations. The dotted line in D and E marks the 0.05 threshold.

The false positive rate and false discovery rate are both very high for the uncorrected procedure (Figure 2BC). *α*-debt and *α*-spending both have a decrease in false positives and false discovery rate when *λ* is positive (Figure 2BC). The false discovery rate for *α*-debt generally lies between the spending (smallest) and investing procedures (largest and one that aims to be below 0.05). Also, for all methods, the true positive rate breaks down as expected when the covariance between variables approaches the noise level. Thus we split the false discovery rate along four quadrants based on *λ* and the noise floor (Figure 2D). The quadrants where true positive covariance is above the noise floor (Q1 and Q2) has a false discovery rate of less than 0.05 for all procedures except uncorrected (Figure 2D). Finally, when varying the number of true positives in the dataset, we found that Q1 and Q2 generally decreases as the number of true positives grow for *α*-spending and *α*-debt, whereas *α*-investing remains the 0.05 mark regardless of the number of true positives (Figure 2E).

All three sequential correction procedures performed well at identifying true positives when these tests were made early on in the analysis sequence. When the “true” tests are later, *α*-debt has the most sensitivity for true positives and *α*-investing is the only procedure that has a stable false discovery rate regardless of the number of true positives (the other two methods appear to be more conservative). The true positive sensitivity and false discovery rate of each of the three sequential correction methods considered depend on the order of statistical tests and how many true positives are in the data.

### Uncorrected sequential tests will flood the scientific literature with false positives

We have demonstrated a possible problem with sequential tests on simulations. We now turn our attention to empirical data from a well known shared dataset in neuroscience to examine the effect of multiple reuses of the dataset. We used 68 cortical thickness estimates from the 1200 subject release of the HCP dataset (Van Essen et al. 2012). We then used 182 behavioural measures ranging from performance during tasks to survey responses (See supplementary table 1) and, for simplicity, ignore all previous publications using the HCP dataset (of which there are now several hundred) for our p-value correction calculation.

We fit 182 linear models in which each behaviour (dependent variable) was modelled as a function of each of the 68 cortical thickness estimates (independent variables), resulting in a total of 12,376 statistical tests. As a baseline, we corrected all statistical tests simultaneously with Bonferroni and FDR. For all other procedures, the independent variables within each mode (i.e. cortical thickness) had simultaneous FDR correction while considering each linear model (i.e. each behaviour) sequentially. The procedures considered were: uncorrected sequential analysis with both Bonferroni and FDR simultaneous correction procedures; all three sequential correction procedures with FDR simultaneous correction within each model. For the sequential tests, the orders were randomized in two ways: (i) uniformly; (ii) weighting the earlier tests to be the significant findings found during the baseline conditions (see Methods). The latter considers how the methods perform if we ask the “right” questions first. Sequential analyses had the order of tests randomized 100 times.

We asked two questions with these models. First, we identified the number of significant findings (p < 0.05, two tail) for the different correction methods. Second, we asked how many additional scientific articles (assuming that at least one positive finding is equal to a publication) would result with the different correction methods. Importantly, in this evaluation of empirical data, we are not necessarily concerned with the number of “true” relationships with this analysis. We care about identifying the number of statistically significant findings within a specified statistical threshold given the different correction procedures. The simultaneous correction procedures act as a baseline. Bonferroni is known to be a conservative procedure. FDR is known to maintain a tolerable ratio of false positives in relation to the number of findings. Thus any procedure that is more stringent than the Bonferroni baseline will be too conservative (more type II errors). Any procedure that is less stringent than FDR will have an increased false discovery rate, implying more false positives (relative to the true positives). Note that, we are tackling only issues regarding correction procedures to multiple hypothesis tests; determining the truth of any particular outcome would require additional replication.

Figure 3 shows the results for all correction procedures. Using sequentially uncorrected tests leads to an increase in significant findings (30/44 Bonferroni/FDR), compared to a baseline of 2 findings when correcting for all tests simultaneously (for both Bonferroni and FDR procedures). Assuming only positive findings are published, this would result in 29/30 (Bonferroni/FDR) publications instead of the baseline 2 (both Bonferroni and FDR), reflecting a 1,400% increase in publications that would primarily reflect false positives.

**Figure 3:**
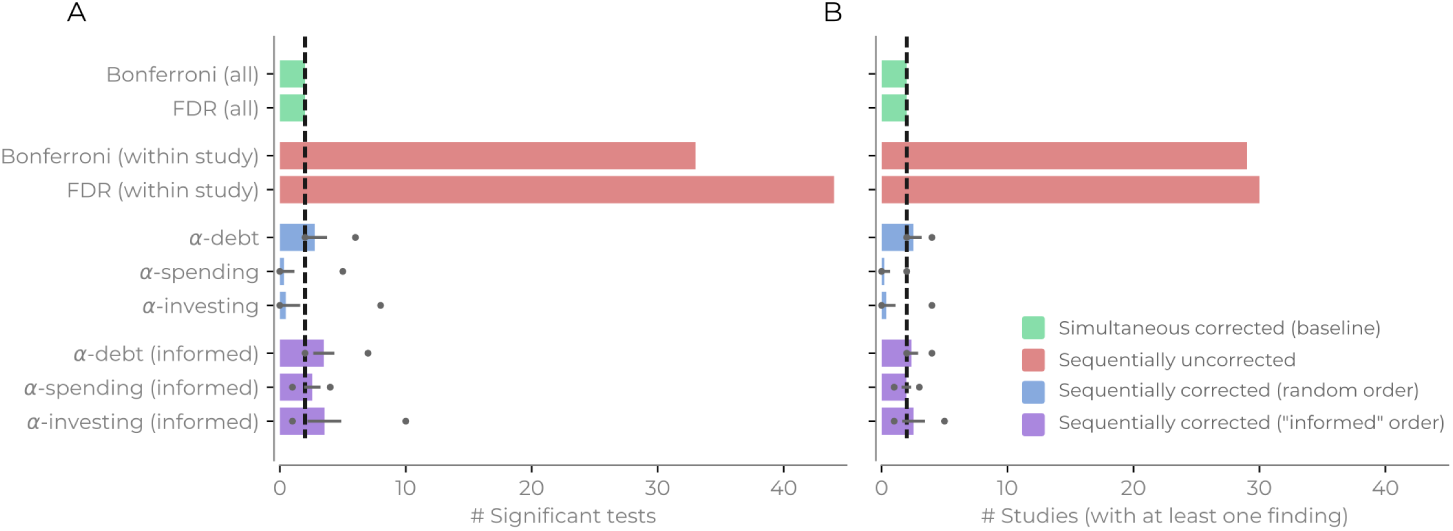
Summary of the number of findings and studies for different correction procedures performed on the empirical dataset. The dotted line shows the baseline form the simultaneous corrections. (A) The number of significant statistical tests for the different correction procedures; (B) Number of studies that had at least one significant finding (and can be considered a possible scientific publication). Error bars show the standard deviation and circles mark min/max number of findings/studies for the sequential correction procedures with a randomly permuted test order.

The sequential correction procedures were closer to baseline but saw divergence based on the order of the statistical tests. If the order was completely random, then *α*-debt found, on average, 2.77 findings (min/max: 2/6) and 2.53 studies (min/max: 2/4) would be published, which is an increase in the number of false positives compared to baseline but considerably less than the sequentially uncorrected procedure. In contrast, *α*-spending found 0.33 (min/max: 0/5) and 0.22 studies (min/max: 0/2) and *α*-investing found 0.48 (min/max: 0/8) findings and 0.37 (min/max 0/4) studies; both of which are below the conservative baseline of 2. When the order is informed by the baseline findings, the sequential corrections procedures increase in the number of findings (findings [min/max]: *α*-debt: 3.49 [2/7], *α*-spending: 2.58 [1/4], *α*-investing: 3.54 [1/10]; studies [min/max]: *α*-debt: 2.38 [2/4], *α*-spending: 1.97 [1/3], *α*-investing: 2.54 [1/5]). All procedures now increase their number of findings above baseline (on average *α*-debt with a random order has a 19% increase in the number of published studies, substantially less than the increase in the number of uncorrected studies). Two conclusions emerge. First, *α*-debt remains sensitive to the number of findings found regardless of the sequence of tests (fewer type II errors) and can never fall above the Bonferroni in regards to type II errors while the other two sequential procedures can be more conservative than Bonferroni. Second, while *α*-debt does not ensure the false positive rate remains under a specific level (more type I errors), it dramatically closes the gap between the uncorrected and simultaneous number of findings and studies.

## Discussion

We have shown with both simulation and an empirical example how sequential statistical tests, if left uncorrected, will lead to a rise of false positive results. Further, we have explored different sequential correction procedures and shown their susceptibility to both false negatives and false positives. Broadly, we conclude that a dataset’s potential to identify new statistically significant relationships will decay over time as the number of sequential statistical tests increases. In the rest of the discussion section we first discuss the implications the different sequential procedures have in regards to the desiderata outlined in the introduction. Then we discuss other possible solutions that could potentially mitigate dataset decay.

### Consequence for sequential tests and open data

We stated three desiderata for open data in the introduction: sharing incentive, open access, and a stable false positive rate. Having demonstrated some properties of sequential correction procedures, we revisit these aims and consider how the implementation of sequential correction procedures in practice would meet these desiderata. The current practice of leaving sequential hypothesis tests uncorrected leads to a dramatic increase in the false positive rate of the scientific literature. While our proposed sequential correction techniques would mitigate this problem, all three require compromising on one or more of the desiderata (summarized in Table 1).

**Table 1 :** Summary of the different sequential correction methods and the open data desiderata. Yes indciates that the method is compatible with the desideratum.

|  | Sharing incentive | Open access | Stable false positive rate |
| --- | --- | --- | --- |
| *α-spending* | No | No | Yes |
| *α-investing* | Yes | No | Yes |
| *α-debt* | Yes | Yes | No |

Implementing *α*-spending would violate the sharing incentive desideratum as it forces the initial analysis to use a smaller statistical threshold to avoid using the entire wealth of *α*. This change could potentially happen with appropriate institutional change, but placing restrictions on the initial investigator (and increased type II error rate) would likely serve as a disincentive for those researchers to share their data. It also places incentives to restrict access to open data (violating the open access desideratum) as performing additional tests would lead to a more rapid decay in the ability to detect true positives in a given dataset.

Implementing *α*-investing, would violate the open access desideratum for two reasons. First, like *α*-spending there is an incentive to restrict incorrect statistical tests due to the sensitivity to order. Second, *α*-investing would require tracking and time-stamping all statistical tests made on the dataset. Given known issues within science, such as the file drawer problem (Rosenthal 1979), this is currently problematic to implement. Also, publication bias for positive outcomes would result in statistical thresholds becoming more lenient over time, which might lead to even more false positives (thus violating no increase in false positives desideratum). Unless all statistical tests are time-stamped, which is possible but would require significant institutional change, this procedure might be hard to implement.

Implementing *α*-debt would improve upon current practices but will compromise on the stable false positive rate desideratum. However, it will have little effect on the sharing incentive desideratum as the original study does not need to account for any future sequential tests. The open access desideratum is also less likely to be compromised as it is less critical to ask the “correct” questions first (i.e. it has the lowest type II error rate of the sequential procedures). However, calculating the correct number of statistical tests performed on a dataset may be practically difficult, given the file drawer problem, and underestimating this number will result in an increased number of false positives. Compared to *α*-investing, estimating the number of tests is a considerably easier task as *α*-debt does not need the order of previous tests and there are conceivable ways of estimating the actual number of sequential tests performed on a dataset. Nevertheless, if a researcher underestimates this number, it will further increase the false positive rate of the method — however will still be better than current practice.

### Other possible procedures

We have only considered frequentist correction procedures to deal with sequential hypothesis testing. There are a few other solutions that are worth exploration, three of which we discuss here. Any of these possible avenues may be superior to the ones we have considered in this article, but they are not readily applicable without some additional consideration.

The first alternative is Bayesian statistics. Multiple comparisons in Bayesian frameworks are often circumnavigated by partial pooling and regularizing priors (Gelman et al. 2013). These techniques should allow for the sequential evaluation of different independent variables against a single dependent variable when using regularizing priors, especially as these different models could also be contrasted explicitly to see which model fits the data best. However, sequential tests could be problematic when the dependent variable changes across experiments. In these multivariate sequential cases, partial pooling cannot be done and regularizing priors may not be sufficient to correct for this. If uncorrected, this could create a similar sequential problem as outlined in this article when inferring relationships between variables in the data. But there are multiple avenues where this could be fixed (e.g. sequentially adjusting the prior odds in Bayes-factor inferences). The extent of sequential analysis on open dataset within the Bayesian hypothesis testing frameworks, and possible solutions, is an avenue of future investigation.

The second alternative is using reusable held-out data. Within machine learning, there have been advances towards having a reusable holdout set in order to query held-out data multiple times (Dwork et al. 2015; Dwork, Hardt, and Roth 2017; Rogers et al. 2019). This avenue is promising, but there appear to be some drawbacks for sequential reuse. First, this line of work within “adaptive data analysis” generally considers a single user querying the holdout test data multiple times while optimizing their model/analysis. Second, this is ultimately a cross-validation technique which is not necessarily the best tool in datasets where sample sizes are small, (Varoquaux 2018) which is often the case with open data and thus not a general solution to this problem. Third, additional assumptions exist in these methods (e.g. there is still a “budget limit” in Dwork et al. (2015) and requires “mostly guessing correctly” in Rogers et al. (2019)). However, this avenue of research has the potential to provide a better solution than what we have proposed here.

The third and perhaps most radical alternative is to reconsider what analyzing open data means. One possible way to handle this problem is to treat all studies using open datasets as a case of exploratory data analysis (EDA), where their primary utility becomes generating hypotheses and testing assumptions of methods (Tukey 1977, 1980; Donoho 2017). Some may consider this reframing problematic, as it could make findings based on open data seem less important. However, accepting that all analysis on open data is EDA would involve less reliance on results from confirmatory statistical inference: the sequential multiple hypothesis test problem disappears. This would lead to an increase of EDA results which may not replicate. However, this is not necessarily problematic; this will not lead to an increase of false positive rate of *confirmatory studies* within the scientific literature but rather would provide a fruitful guide about which confirmatory studies to undertake. Those who consider open data’s value to be more than exploratory will naturally disagree with this perspective. Implementing this suggestion would require little infrastructural or methodological change; however, it would require an institutional shift in how researchers interpret open data results.

### Conclusion

One of the benefits of open data is that it allows multiple perspectives to approach a question, given a particular sample. The trade-off of this benefit is that more false positives will enter the scientific literature. We remain strong advocates of open data and data sharing, but researchers using openly shared data must be sensitive to the accumulation of false positives and ensuing dataset decay that will occur with repeated reuse. Ensuring findings are replicated using independent samples will greatly decrease the false positive rate, since the chance of two identical false positives relationships occurring, even on well explored datasets, is small.

## Methods

### Simulations

The first simulation sampled data for one dependent variable and 100 independent variables from a multivariate Gaussian distribution (mean: 0, standard deviation: 1, covariance: 0). We conducted 100 different pairwise sequential analyses in a random order. For each analysis, we quantified the relationship between an independent variable and the dependent using a Pearson correlation. If the correlation had a two-tailed p-value less than 0.05, we considered it to be a false positive. The simulation was repeated for 1,000 iterations. The probability of at least one false positive is the average number of iterations where there was at least one false positive analysis.

The second simulation had three additional variables. First, a variable that controlled the number of true positives in the data. This variable varied between 1-10. Second, the selected true positive variables, along with the dependent variable, had their covariance assigned as *p*. *p* varied between 0 and 1 in steps of 0.025. Finally, we wanted to test the effect of the analysis order to identify true positive variables. Each sequential analysis, (*m*_1_, *m*_2_, *m*_3_ …), could be assigned to be a “true positive” (i.e. covariance of *p* with the dependent variable) or a “true negative” (covariance of 0 with dependent variable). First, *m*_1_ would be assigned one of the trials, then *m*_2_ and so forth. This procedure continued until there were only true positives or true negatives remaining. The procedure assigns the *i*th analysis to be randomly assigned, weighted by *λ*. If *λ* was 0, then there was a 50% chance that *m*_*i*_ would be a true positive or true negative. If *λ* was 1, a true positive was 100% more likely to be assigned to *m*_*i*_ (i.e. an odds ratio of 1+*λ*:1), The reverse occurred if *λ* was negative (i.e. -1 meant a true negative was 100% more likely at *m*_*i*_).

### Empirical example

Data from the Human Connectome Project (HCP) 1200 subject release was used (Van Essen et al. 2012). We selected 68 estimates of cortical thickness to be the indepndent variables for 182 continuous behavioural and psychological variables dependent variables. Whenever possible, the age-adjusted values were used. Supplementary Table 1 shows the variables selected in the analysis.

For each analysis, we fit an ordinary least squares model was fit using statsmodels (0.10.0-dev0+1579, https://github.com/statsmodels/statsmodels/). For all statistical models, we first standardized all variables to have a mean of 0 and a standard deviation of 1. We dropped any missing values for a subject for that specific analysis. Significance was considered for any independent variable if it had a p-value < 0.05, two-tailed for the different correction methods.

We then quantified the number of findings and the number of potential published studies that the different correction methods would present. The number of findings is the sum of significant independent variables. The number of potential studies is the number of dependent variables that had at least one significant finding. The rationale for the second metric is to consider how many potential findings would exist in the literature if a separate group conducted each analysis, and only significant findings were published.

For the sequential correction procedures, we used two different sequential tests orderings. The first was with a uniformly random order. The second was an “informed” order that pretends we somehow *a priori* knew which variables will be correlated. The “informed” order was created by identifying the significant statistical tests when using simultaneous correction procedures (FDR and Bonferroni, see below) which acted as a baseline to identify analyses which were considered “baseline positives” (i.e. significant with simultaneous FDR. These were two analyses) and the other analyses that were “baseline negatives”. Then, as in simulation 2, the first analysis *m*_1_ was randomly assigned to be positive or negative with equal probability. This “informed” ordering means that the “baseline positives” would usually appear in an earlier order than when the sequence order was sequentially randomized. All sequential correction procedures were applied 100 times with the sequence order randomized.

### Simultaneous correction procedures

We used the Bonferroni method and the Benjamini & Hochberg FDR method for simultaneous correction procedures (Benjamini and Hochberg 1995). Both correction methods were run using multipy (v0.16, https://github.com/puolival/multipy). In the simulations over multiple iterations, the false discovery rate calculation was based on the average false positives and the average true positives over the iterations.

### Sequential correction procedures

#### Uncorrected

This procedure is to not correct for any sequential analysis. This analogous to reusing open data with no consideration for any sequential tests that occur due to data reuse. For all sequential hypothesis tests, p<0.05 signified statistical significance.

#### α-debt

A sequential correction procedure that, to our knowledge, has not previously been proposed. At the first hypothesis tested, *α*_1_ sets the statistical significance threshold (here 0.05). At the *i*th hypothesis tested the statistical threshold is 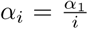. The rationale here is that at the *i*th test, a Bonferroni correction is applied that considers there to be *i* number of tests performed. This method lets the false positive rate increase (i.e. the debt of reusing the dataset) as each test corrects for the overall number of tests, but all earlier tests have a more liberal threshold. The total possible “debt” incurred for *m* number of sequential tests can be calculated by 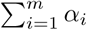 and will determine the actual false positive rate.

#### α-spending

A predefined *α*_0_ is selected which is called the *α*-wealth. At the *i*th test the statistical threshold, *α*_*i*_, a value is selected to meet the condition that 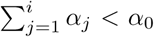. The *i*th test select *α*_*i*_ that spends part of the remaining “*α*-wealth”. The remaining *α*-wealth at test *i* is 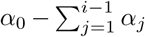. Like, *α*-debt, this method effectively decreases the p-value threshold of statistical significance at each test. However, it can also ensure that the false positive rate of all statistical tests is never higher than *α*_0_. Here, at test *i* we always spend 50% of *α*_*i*−1_ and *α*_0_ is set to 0.05. See (Foster and Stine 2008) for more details.

#### α-investing

The two previous methods only allow for the statistical threshold to decrease over time and are more akin to familywise error correction procedures. An alternative approach, which is closer to false discovery rate procedures, is to ensure the false discovery rate remains below a predefined wealth value (*W*_0_) (Foster and Stine 2008). At each test, *α*_*i*_ is selected from the remaining wealth at *W*_*I* − 1_. If the sequentially indexed test *i* was considered “statistically significant” (i.e. rejecting the null hypothesis), then *W*_*i*_ increases: *W*_*i*_ = *W*_*i*−1_ +*ω*. Alternatively, if the null hypothesis cannot be rejected at *i*, then the wealth decreases: 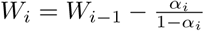. We set *ω* to *α*_0_, which is the convention, *α*_0_ to 0.05, and *α*_*i*_ is set to 50% *-*of the remaining wealth. See (Foster and Stine 2008) for more details.

When combining the simultaneous and sequential correction procedures in the empirical example, we used the sequential correction procedure to derive *α*_*i*_, which we then used as the threshold in the simultaneous correction.

### Data/Code availability statement

Code for the simulations and analyses is available at https://github.com/wiheto/datasetdecay. The data is openly available at the Human Connectome Project at https://db.humanconnectome.org.

## Acknowledgements

WHT acknowledges support from the Knut och Alice Wallenbergs Stiftelse (SE) (grant no. 2016.0473, http://kaw.wallenberg.org). We would also like to thank Pontus Plavén-Sigray, Lieke de Boer, Nina Becker, Granville Matheson, Björn Schiffler, and Gitanjali Bhattacharjee for helpful discussions and feedback.

## Supplementary Materials

### Sequential family examples

Recall that there are two definitions when tests are part of the same family: (i) to prevent data-dredging, and (ii) for “confirmatory” analyses when tests are a “conceptual family” by supporting “similar research questions” (Hancock and Klockars 1996). Recall also that our rule-of-thumb that sequential tests are part of the same family if they would be considered part of the same family in a simultaneous test.

We have to consider that all data reuse could be considered a type of data dredging as, for some, pre-specifying a hypothesis is always before data collection (Tukey 1991). If we consider a study that, after some initial analyses, makes a new hypothesis and analyses this, this would be considered secondary, *post hoc*, or data dredging. The only difference in data reuse is that a different researcher has performed the analysis/hypothesis. Some may argue that confirmatory work is possible with open data, especially if the research question is different enough (see an example of separated families below). However, others may argue that, especially if the researchers know of a paper using that data, it should be exploratory. Thus, it appears that many cases of data reuse should fall within the exploratory category.

However, if a researcher can justify that their analysis is “confirmatory” research, the next question is whether they are helping to answer “associated research question” as previous research. This answer is not always clearcut and can be challenging to determine. There are some clear cut cases and examples where it is not always apparent when the hypothesis is considered confirmatory:

#### A clear example of the same family

In the empirical demonstration in the main text, we test multiple personality and behavioural variables. If all these tests were considered confirmatory, there is substantial overlap in the research question here which ultimately boils down to what can cortical thickness explain.

#### A clear example of separated families

Many datasets have variables that will be used in most analyses using that dataset, but this is not always the case. The PubMed Central Open Access Subset dataset contains hundreds of thousands of academic articles. This dataset can be used in many ways different ways to approach multiple very different research questions stretching from the gender of authors, readability of writing, or the semantic similarity of research topics. These are all very different research questions and become different families. If presented in the same article, there is little to no overlap of the research questions being part similar; thus, these are different families.

#### Gray-area when families are not clear

Consider an fMRI dataset analysed in multiple different ways (e.g. a mass univariate analysis or a connectivity analysis that transforms voxel time series into an adjacency matrix). Let us assume these two different ways of representing fMRI data correlate with the same variable (task performance). Are these the same research questions? It depends, (Hochberg and Tamhane 1987) noted that different families could share statistical dependence with each other, which is the case here, so they are not necessarily the same family just because the data shares some statistical relationship. The question boils down to if they are answering the same research question or not. If both analyses are trying to explain task performance, they are the same family as it is a similar research question. Whereas if the research is about how to quantify brain data (as a network or as regions), it can be argued that they are different families.

It seems pertinent that researchers must at least reflect on why their sequential tests are not part of the same family as previous tests if they decide to not correct for them. This reflection should: (i) justify that their study meets the requirements for being confirmatory, and (ii) justify how their particular set of tests should be considered a different family of tests from any previous studies using the data. The second will depends on whether the dataset has a variable that most analyses will use in a similar way to tackle a similar set of research questions. For example, polling data will use “voting intention” in most analyses, and neuroimaging datasets will generally study the same cognitive process/task (and not ask questions about representations). In many instances, these two criteria will be hard to meet.

